# Revealing allostery in *PTPN11* SH2 domains from MD simulations

**DOI:** 10.1101/2022.06.29.498118

**Authors:** Massimiliano Anselmi, Jochen S. Hub

## Abstract

Src-homology 2 (SH2) domains are protein interaction domains that bind to specific peptide motifs containing phosphotyrosine. SHP2, a tyrosine phosphatase encoded by *PTPN11* gene, which has been emerged as positive or negative modulator in multiple signaling pathways, contains two SH2 domains, respectively called N-SH2 and C-SH2. These domains play a relevant role in regulating SHP2 activity, either by recognizing its binding partners or by blocking its catalytic site. Considering the multiple functions that these domains carry out in SHP2, N-SH2 and C-SH2 represent an interesting case of study. Here, we present a methodology that permits, by means of the principal component analysis (PCA), to study and to rationalize the structures adopted by the SH2 domains, in terms of the conformations of their binding sites. The structures can be distinguished, grouped, classified and reported in a diagram. This approach permits to identify the accessible conformations of the SH2 domains in different binding conditions and to eventually reveal allosteric interactions. The method further reveals that the conformation dynamics of N-SH2 and C- SH2 strongly differ, which likely reflects their distinct functional roles.

## 1. Introduction

A cell needs to continuously respond to external and internal stimuli to perform its processes. For this purpose, the cell response is often mediated by covalent modifications of existing proteins. Several posttranslational modifications are found in nature, including phosphorylation and dephosphorylation of several amino acid residues, such as serine, threonine and tyrosine. Posttranslational modifications are recognized and transduced into signal cascades via protein interaction domains, which often recognize short peptide motifs of their target protein. The interaction domains usually contain a conserved binding site with high affinity for the modified residue, along with a more variable cleft that, engaging the flanking residues, confers selectivity and allows to distinguish different peptide motifs containing the same posttranslational modification [1]. Protein tyrosine phosphorylation (PTP) is an important posttranslational modification that controls the cell signaling involved in a variety of biological processes, including cell growth, proliferation, differentiation, migration, survival, and apoptosis.

Src-homology 2 (SH2) domains are protein interaction domains of about 100 amino acids that bind to specific peptide motifs containing phosphotyrosine (pY). Atomic structures of a large number of SH2 domains have been determined. SH2 domains share a common fold consisting of a central anti-parallel β-sheet flanked by two α-helices (Figure 1). Phosphopeptides bind in extended conformation along a cleft that lies perpendicular to the central β-sheet. Conserved residues contribute to the hydrophobic core or are involved in pY recognition, while more variable residues contribute to specific recognition of the residues C-terminal to the pY [2].

**Figure 1.**
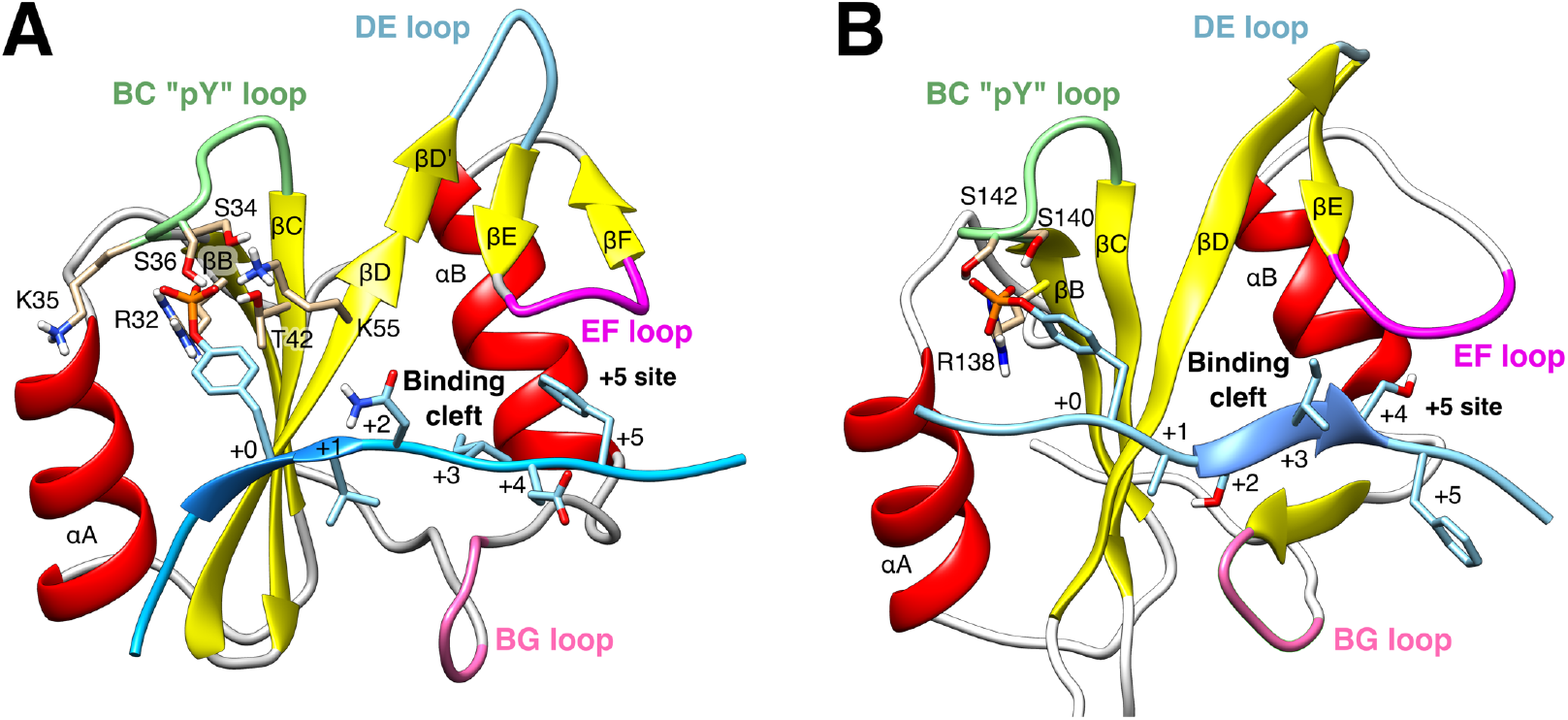
Cartoon representations (A) of the N-SH2 domain in complex with the SPGEpYVNIEFGS (IRS-1 pY895) peptide (crystal structure, PDB ID 1AYB) [15] and (B) of the C-SH2 domain in complex with the LSApYASISFQK (IRS-1 pY1229) peptide, taken from an MD simulation. The peptide is shown in cyan, comprising the phosphotyrosine and residues from position +1 to +5 relative to the phosphotyrosine (see labels). Functionally important loops are highlighted in color: BC “pY” loop (green), DE loop (light blue), EF loop (magenta), BG loop (deep pink). The phosphotyrosine binds the site delimited by the pY loop and the central β-sheet (βB, βC, βD strands). EF and BG loops delimit the “+5 site” where the peptide residue in position +5 is settled.

Amongst all SH2 domains, those belonging to SHP2, a tyrosine phosphatase encoded by *PTPN11* gene, represent a case of study with major physiological and medical impact. SHP2 is a regulator of signaling downstream of several cell surface receptors, functioning as positive or negative modulator in multiple signaling pathways. The structure of SHP2 includes two tandemly-arranged SH2 domains, called N-SH2 (Figure 1A) and C-SH2 (Figure 1B), followed by the catalytic PTP domain, and a C-terminal tail. Under basal conditions, SHP2 is in an autoinhibited state where the N-SH2 domain occludes the catalytic site of the PTP domain. Association of SHP2 to its binding partners via the SH2 domains favors the release of the autoinhibitory N-SH2–PTP interactions, rendering the catalytic site accessible to substrates [3].

Thus, N-SH2 plays the two-fold role *(i)* of blocking the catalytic site in the autoinhibited structure and *(ii)* of recognizing the binding partners before the activation takes place. For that reason, a coupling is required between the binding of a phosphopeptide to N-SH2 and the loss of affinity of N-SH2 for the PTP domain, which is likely realized via a series of conformational changes. On the other hand, C-SH2 recognizes one of the two phosphorylated motifs in BTAMs, namely a kind of ubiquitous phosphorylated bisphosphoryl tyrosine-based activation motifs. BTAMs, which contain two phosphorylated motifs separated by a linker, activate SHP2 by binding to both N-SH2 and C- SH2. The second recognition site in C-SH2 confers a grade of conformational selection that favors, in presence of a linker of suitable length, the complete displacement of the N-SH2 domain from the PTP active site [3]. Hence, C-SH2 requires a pY recognition site for its function similar to N-SH2, but it may not require the allosteric mechanism that triggers the release of N-SH2 from the PTP domain upon the ligand binding.

In this chapter, we present a simple and effective methodology that permits, by means of the principal component analysis (PCA) [4], to study and to rationalize the structures adopted by the SH2 domains, in their apo and holo form. The conformational states are reported in an essential plane defined by two eigenvectors, respectively describing the principal mode either of the affinity site or of the specificity site. The structures can be distinguished, grouped and classified in terms of the conformations adopted by the affinity and specificity sites. This approach permits to report the conformational space explored by a SH2 domain in different binding conditions, and to quickly compare conformational distributions. It is also possible to highlight the allosteric coupling between the affinity and specificity sites in a SH2 domain, or to compare the conformations adopted by two distinct SH2 domains, if the aminoacidic sequence is sufficiently conserved. Accordingly, a comparison between the N-SH2 domain and the C-SH2 domain of SHP2 suggests that the conformational rearrangement, put in place to accommodate a phosphopeptide, is more marked in N-SH2 than in C-SH2. We hypothesize that this difference is linked to the peculiar role of N-SH2 in SHP2 activation, namely to recognize a binding partner and, at the same time, to control the phosphatase activity of SHP2.

## 2. Materials

Molecular dynamics simulations were performed on the isolated N-SH2 and C-SH2 domains in solution, complexed with a set of high affinity phosphopeptides, with natural or synthetic sequence. SLNpYIDLDLVK (IRS-1 pY1172) [6], IEEpYTEMMPAA (IRS-1 pY546) [7], QVEpYLDLDLD (Gab1 pY627) [8], SVLpYTAVQPNE (PDGFR-β pY1009) [7], AALNpYAQLMFP [9], RLNpYAQLWHR [9], SPGEpYVNIEFGS (IRS-1 pY895) [7], VLpYMQPLNGRK [10] were complexed with the N-SH2 domain (Table 1), whereas HTEpYASIQTSP (SHPs-1 pY453) [11], FSEpYASVQVPRK (SHPs-1 pY496) [11], LSApYASISFQK (IRS-1 pY1229) [12], RVDpYVVVDQQK (Gab1 pY659) [13], TMVpYGHLLQDS (CDCP1 pY743) [14] were complexed with the C-SH2 domain (Table 2). The initial atomic coordinates of N-SH2 domains were taken from crystallographic structures [15,16]. The peptides bound to the N-SH2 domain were modeled by means of Molecular Operative Environment (MOE) [17] (Table 1). The structures of the C-SH2 domain bound to phosphopeptides were obtained by homology from the crystal structure of PTPN11 tandem SH2 domains in complex with TXNIP peptide (PDB ID 5df6) [18]. The peptides were docked to the C-SH2 binding cleft by CLUSPRO [19] (Table 2). N-SH2 domains comprised the residues from position 3 to 103. C-SH2 domains comprised the residues from position 109 to 217. All MD simulations were performed with the GROMACS software package [20], using AMBER99SB force field [21]. Production simulations of 1 µs were performed. More details on the setup of MD simulations can be found in our previous publication [22].

**Table 1.**
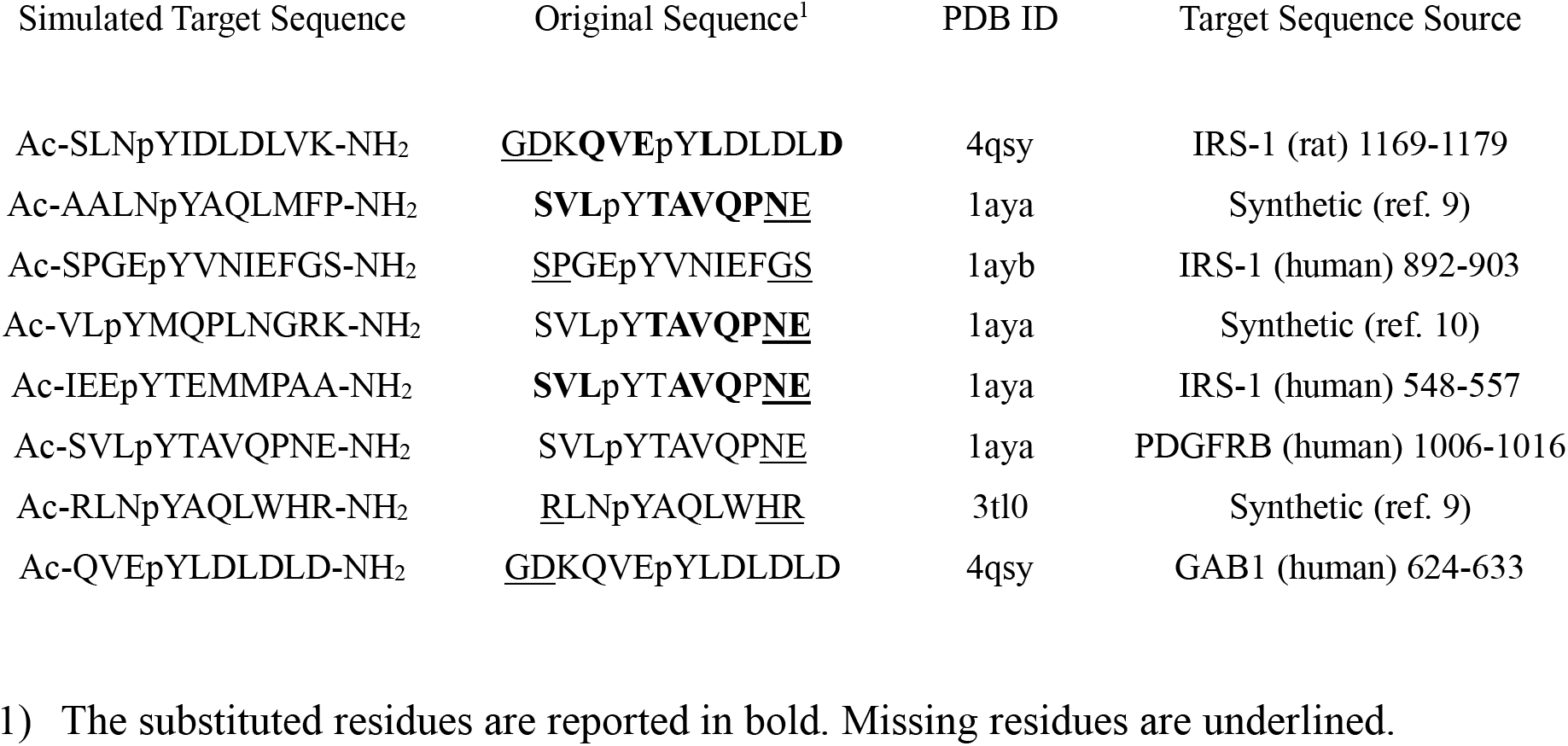
Peptides bound to the N-SH2 domain.

**Table 2.**
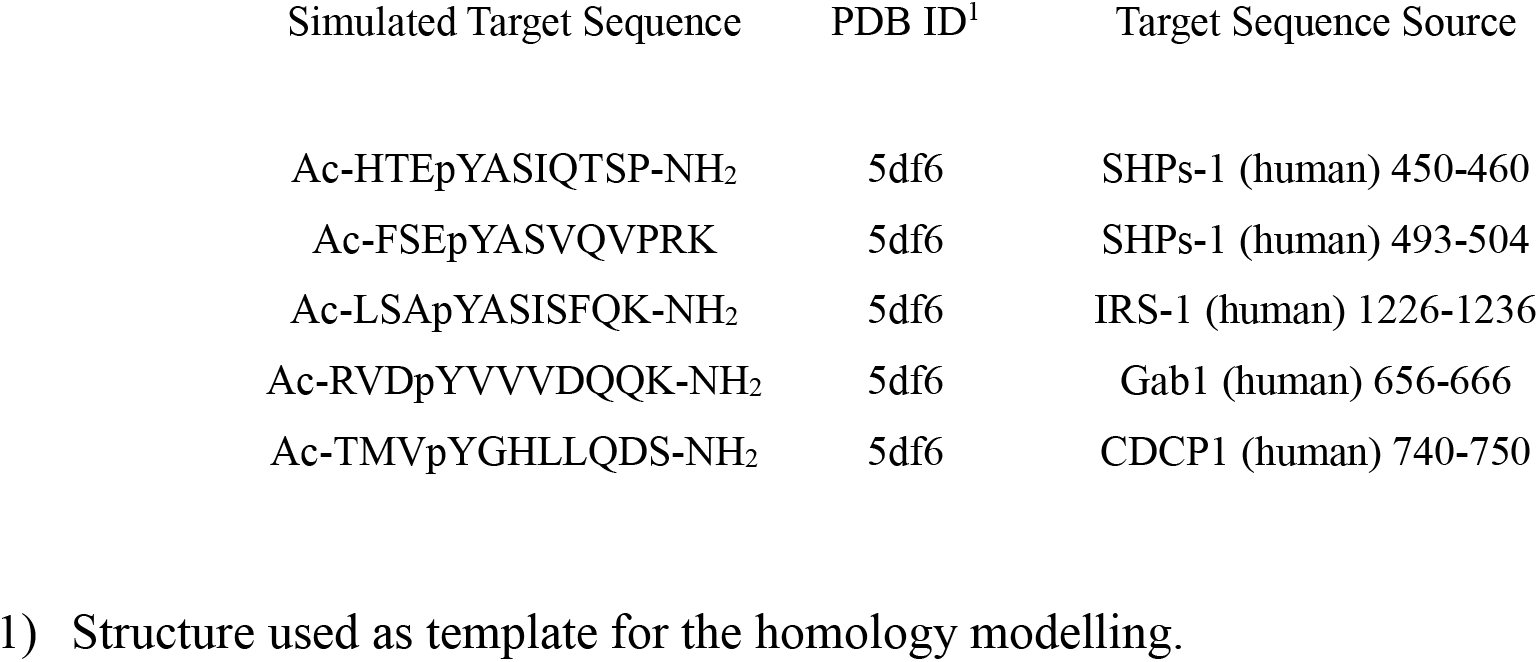
Peptides bound to the C-SH2 domain.

| Simulated Target Sequence | PDB ID<sup>1</sup> | Target Sequence Source |
| --- | --- | --- |
| Ac-HTEpYASIQTSP-NH<sub>2</sub> | 5df6 | SHPs-1 (human) 450-460 |
| Ac-FSEpYASVQVPRK | 5df6 | SHPs-1 (human) 493-504 |
| Ac-LSApYASISFQK-NH<sub>2</sub> | 5df6 | IRS-1 (human) 1226-1236 |
| Ac-RVDpYVVVDQQK-NH<sub>2</sub> | 5df6 | Gab1 (human) 656-666 |
| Ac-TMVpYGHLLQDS-NH<sub>2</sub> | 5df6 | CDCP1 (human) 740-750 |

1) Structure used as template for the homology modelling.

## 3. Methods

### 3.1 Theory

Identifying the motions of interest in a biomolecule, in order to uncover functional mechanisms, is a non-trivial task considering the complexity of the system. In molecular dynamics (MD) simulations, both the rapid local fluctuations and the slow collective motions occur simultaneously and in erratic manner, making it hard to distinguish functionally relevant motions from thermal noise. Covariance analysis, also called principal component analysis (PCA), is a useful tool to detect correlated motions in protein dynamics. The PCA assumes that the large-scale collective modes of motion dominate the functional dynamics, so that the majority of protein dynamics can be described by a low number of collective degrees of freedom. This approach has the advantage to identify the main modes of collective motions and to filter them from the noisier local fluctuations. Moreover, it is possible to easily describe complex structural rearrangement in the high-dimensional conformational space by means of a few collective variables [5].

After superposition to a common reference structure, a non-mass weighted covariance matrix ***C*** of the atomic coordinates of *N* atoms is constructed:

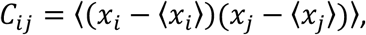

where ⟨. ⟩ denotes the average over all simulation frames. ***C*** is a symmetric 3*N*×3*N* matrix that can be diagonalized by an orthogonal coordinate transformation ***R***:

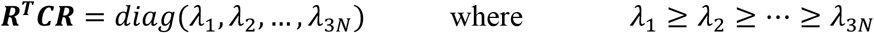

The eigenvalues λ_*i*_ correspond to the mean square fluctuation along the *i*^th^ eigenvector, and, therefore, represent the contribution of each principal component to the total fluctuations. The eigenvectors are sorted such that their eigenvalues are in decreasing order. Usually, six eigenvalues are exactly zero, since the corresponding eigenvectors describe the overall rotation and translation that are eliminated by the superposition. The columns of the orthogonal matrix ***R*** are the eigenvectors, also called principal or essential modes. Let ***ξ***_***i***_ denote the *i*^th^ eigenvector of ***C*** (the *i*^th^ column of ***R***), then the projection of the MD trajectory **x**(*t*) onto the *i*^th^ principal mode gives the principal component *p*_*i*_:

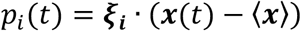

Finally, the trajectory can be filtered along one (or several) principal modes, thereby transforming the projections onto the eigenvectors back to Cartesian coordinates:

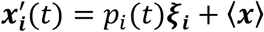

### 3.2 PCA of the cumulative trajectory

To characterize the conformations that the N-SH2 domain may adopt, apply principal component analysis (PCA) to the cumulative trajectory from all simulations of the N-SH2 domain bound to different phosphopeptides (Table 1) [22]. We assume that the simulation trajectories were previously converted so that the domain is whole in all frames. Since the phosphopeptides differ and the cumulative trajectory can include only a fixed number of ordered atom coordinates, consider only the part of the system shared among all simulations, namely the N-SH2 domain. To this end, use the *trjconv* and *trjcat* modules of GROMACS to extract the N-SH2 domain coordinates from each trajectory and to concatenate the coordinates into a single cumulative trajectory:

~~~
$ gmx trjconv -f traj_1.xtc -n index_1.ndx -o traj_N-SH2_only_1.xtc
$ gmx trjconv -f traj_2.xtc -n index_2.ndx -o traj_N-SH2_only_2.xtc
…
$ gmx trjcat -f traj_N-SH2_only_*.xtc -o cumulative_traj_N-SH2_only.xtc -settime
~~~

Then calculate and diagonalize the non-mass weighted covariance matrix by the *covar* module:

~~~
$ gmx covar -f cumulative_traj_N-SH2_only.xtc -s ref_structure_N-SH2.gro \
  -n index.ndx
~~~

Build the covariance matrix using the backbone atoms from residues Trp^6^ to Pro^101^, excluding the flexible termini. Superimpose the structures by a least-squares fit onto the backbone considering only the residues with small root mean-squared fluctuation, representing the relatively rigid core of the domain. The core is defined by the sequence ranges Phe^7^–Pro^33^, Asp^40^–Arg^47^, Ala^50^–Asn^58^, Asp^61^–Leu^65^, Phe^71^–Tyr^81^, Leu^88^–Glu^90^, Val^95^–Pro^101^. It is worth noting that the PCA eigenvectors do not necessarily represent the conformational transitions undergone by the N-SH2 domain during an individual simulation; instead, they describe the overall conformational space accessible to the N-SH2 domain, as collected from all simulations (see Note 1). Use the output file *eigenvec*.*trr* containing the eigenvectors for the following analysis.

Analyze the eigenvectors by the *anaeig* module:

~~~
$ gmx anaeig -f cumulative_traj_N-SH2_only.xtc -s ref_structure_N-SH2.gro \
  -v eigenvec.trr -first 1 -last 1 -proj proj.xvg -extr extreme.pdb \
  -nframes 30
~~~

In this example, only the first eigenvector is analyzed (options *-first* and *-last*), the trajectory of the projection onto the first eigenvector is saved in *proj*.*xvg* while a trajectory of 30 frames joining the extreme configurations along the first eigenvector is saved in *extreme*.*pdb*.

### 3.3 Large-scale correlated motion revealed by standard PCA

PCA suggests that the N-SH2 domain may adopt at least two main conformational states, hereafter called α and β [22]. In this example, these two states are resolved by the first PCA vector, which is representative of almost 40% of the overall domain fluctuations. The correlated structural rearrangements detected by the first PCA vector are visualized in Fig. 2A by means of the extreme projections onto the vector (see also Fig. 3A), and the related structural rearrangements are quantified in Fig. 2E–G (see Note 2). Namely, the α-state shown in Fig. 2A (transparent) is characterized by (i) a closed pY loop (Lys^35^ Cα–Thr^42^ Cα distance ∼ 9 Å), (ii) an increased distance between the ends of two β-strands, βC and βD, leading to breaking of three inter-strand hydrogen bonds and spreading of the central β-sheet into a Y-shaped structure (Gly^39^ C–Asn^58^ N distance ∼ 12 Å), and (iii) the closed +5 site with a narrow, less accessible cleft (Tyr^66^ Cα–Leu^88^ Cα distance ∼ 7 Å; see also Fig. 3A, left). The β-state shown in Fig. 2A (opaque) is characterized by (i) an open pY loop (Lys^35^ Cα–Thr^42^ Cα distance ∼ 11 Å), (ii) a closed central β-sheet with parallel β-strands (Gly^39^ C–Asn^58^ N distance ∼ 4 Å), and (iii) an open +5 site with an accessible cleft (Tyr^66^ Cα–Leu^88^ Cα distance ∼ 12 Å, see also Fig. 3A, right). The correlation analysis shows that the conformation adopted by the affinity binding site (pY loop) is strictly coupled to the spread of the central β-sheet (Fig. 2B, correlation R = −0.82) and strongly associated with the opening/closure of the specificity binding site (+5 site) (Fig. 2D, R = 0.64) [22].

**Figure 2.**
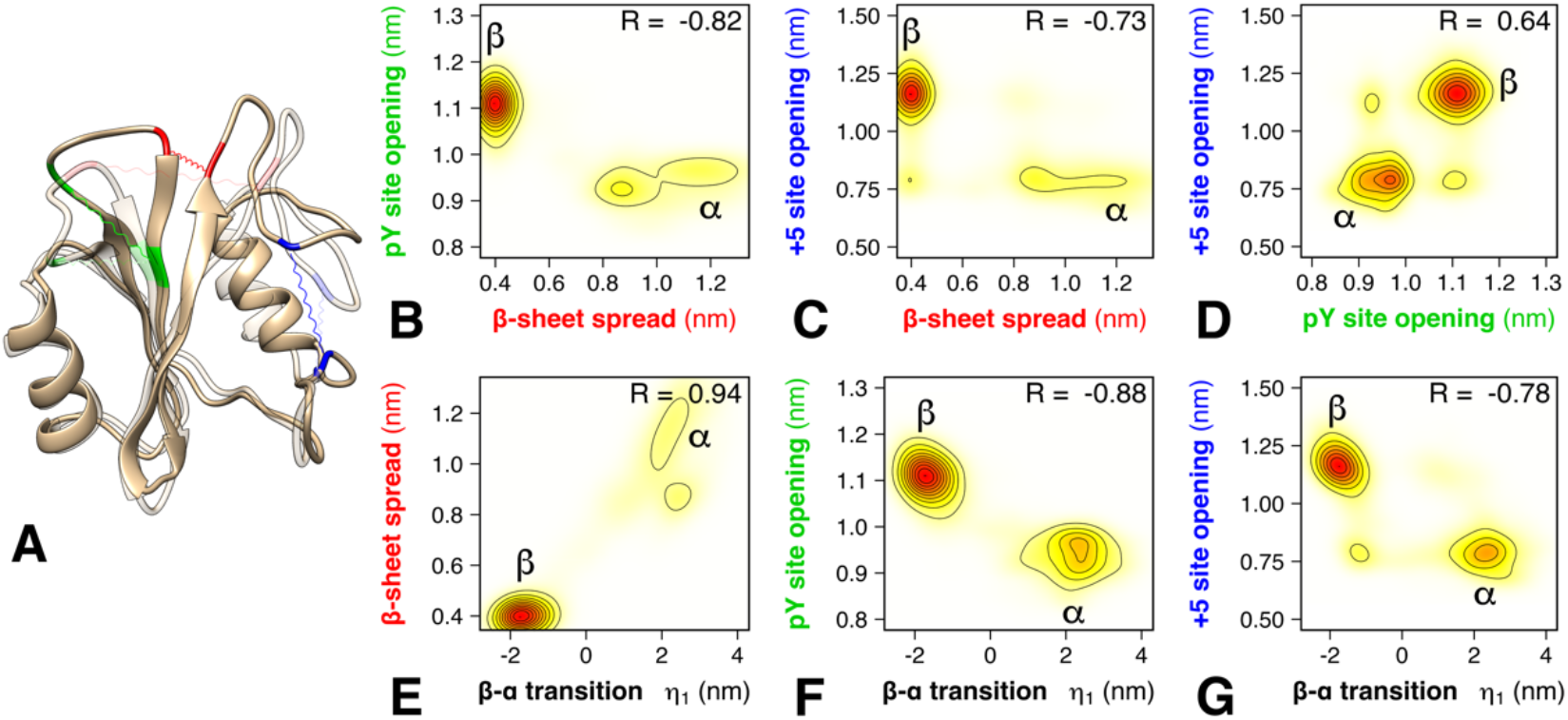
Conformational states and correlations revealed by the first PCA vector. (A) Conformational transition from β (opaque) to α (transparent), representing the two main conformational states adopted by the N-SH2 domain, here visualized as the extreme projections onto the first PCA vector. The residues used to quantify the β-sheet spread, the pY loop opening, and the +5 site opening are highlighted in red, green, and blue, respectively. (B–D) Correlation between the β-sheet spread, pY loop opening, and +5 site opening, as taken from microsecond simulations of N-SH2 bound to 8 different peptides. The distances were defined as described in the main text. (E–G) Correlation between the projection η_1_ onto the first PCA vector and β-sheet spread, pY loop opening, and + 5 site opening (see axis labels). Pearson correlation coefficients R are shown in each panel (B–G). Reproduced from Scientific Reports 2020 with permission from Springer Nature [22].

**Figure 3.**
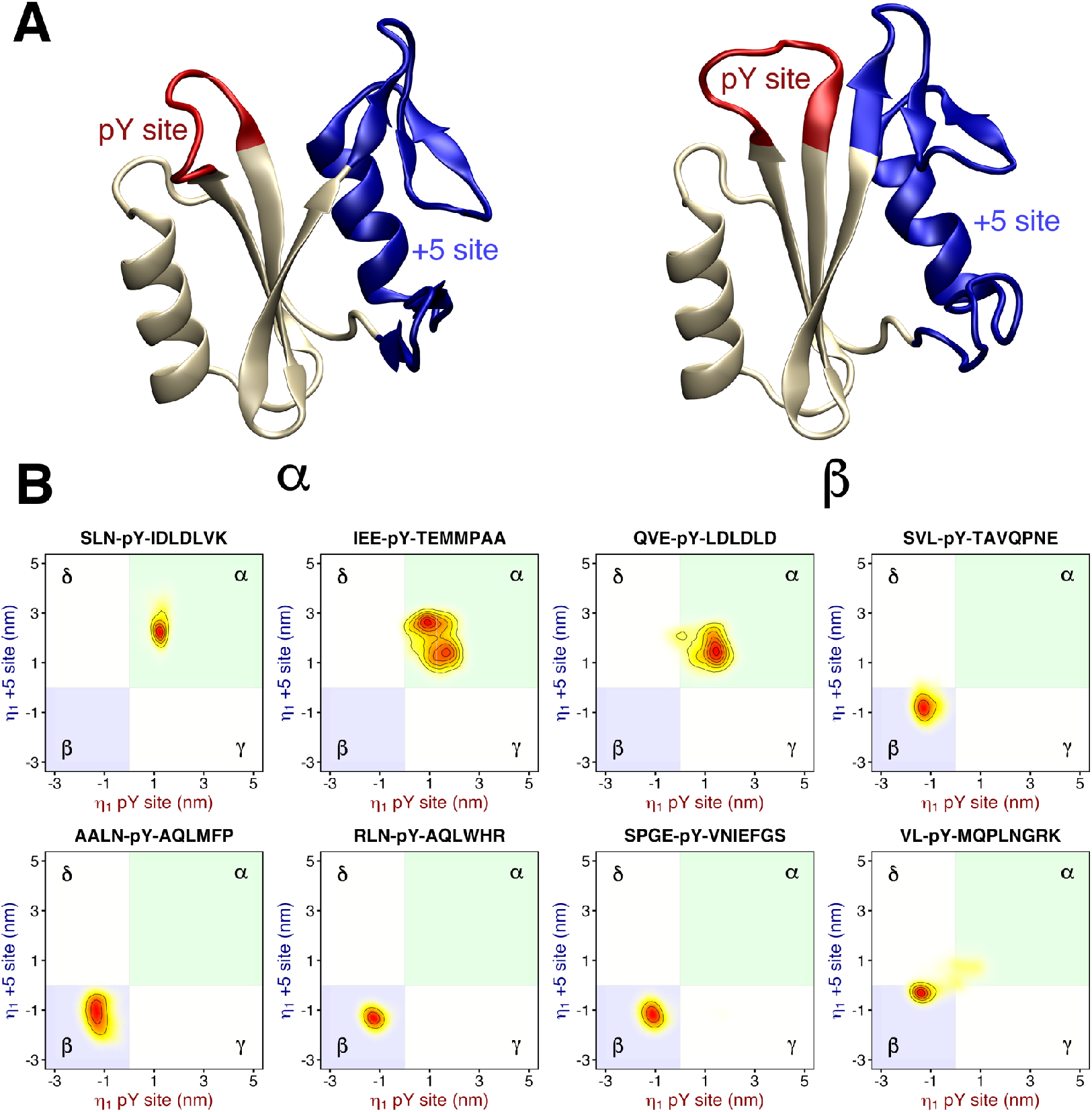
(A) Cartoon representation of the two conformations, α (left) and β (right), adopted by the N-SH2 domain, and shown as the extreme projections onto the first PCA vector. From the first PCA vector, two subsectors were selected, describing the motions of the pY loop (Ser^34^–Phe^41^, red cartoon) and of the +5 site (Gln^57^–Glu^97^, blue cartoon), respectively. The residues used for defining the +5 site comprise the end of the βD strand, the DE loop, the EF loop, the αB helix and the BG loop. (B) Projection of N-SH2 trajectories with different bound peptides (subplot titles) projected onto the PCA subvectors of the pY loop (abscissa) and of the +5 site (ordinate). The region corresponding to the α-state (pY loop closed, +5 site closed) is shaded in green, the region of the β- state (pY loop open, +5 site open) is shaded in blue. The regions corresponding to the additional γ- state (pY loop closed, +5 site open) and δ-state (pY loop open, +5 site closed) are reported as white areas. Reproduced from Scientific Reports 2020 with permission from Springer Nature [22].

### 3.4 Allosteric coupling highlighted by decomposition of the PCA vector into structural motifs

The covariance analysis on the cumulative trajectory suggests a coupling between the pY loop and the +5 site (see Note 3), revealing an allosteric interaction for controlling the state of the pY loop through binding of a specific amino acid sequences at the +5 site (Fig. 2D). To analyze the coupling during simulations with different peptides, split the PCA vector into two subvectors (comprising Ser^34^–Phe^41^ and Gln^57^–Glu^97^ backbone respectively) that describe the motion either of the pY loop or of the +5 site.

Technically, these subvectors are generated by PCA considering only the respective subsets of atoms in a filtered trajectory which has been previously projected onto the full-length PCA vector *ξ*_***1***_. At this purpose, use the option *-filter* of the *anaeig* module, and then calculate and diagonalize the covariance matrix with the *covar* module for only the residues Ser^34^–Phe^41^ or Gln^57^–Glu^97^ (see Note 4):

~~~
$ gmx anaeig -f cumulative_traj_N-SH2_only.xtc -s ref_structure_N-SH2.gro   \
  -v eigenvec.trr -first 1 -last 1 -filter cumulative_traj_filtered.xtc
$ gmx covar -f cumulative_traj_filtered.xtc -s ref_structure_N-SH2.gro   \
  -n index.ndx -o eigenvec_pYsite.trr
$ gmx covar -f cumulative_traj_filtered.xtc -s ref_structure_N-SH2.gro   \
  -n index.ndx -o eigenvec_+5site.trr
$ gmx anaeig -v eigenvec_pYsite.trr -f traj_N-SH2_only_1.xtc -n index.ndx   \
  -s ref_structure_N-SH2.gro -first 1 -last 1 -proj proj_pYsite_1.xvg
$ gmx anaeig -v eigenvec_+5site.trr -f traj_N-SH2_only_1.xtc -n index.ndx   \
  -s ref_structure_N-SH2.gro -first 1 -last 1 -proj proj_+5site_1.xvg
~~~

Finally, plot the projection onto the pY site subvector versus the projection onto the +5 site subvector using a combination of Unix commands, sed, awk, and paste:

~~~
$ paste <(sed ′/^[#@&]/d’ proj_pYsite_1.xvg | awk ′{print $2}’) \
        <(sed ′/^[#@&]/d’ proj_+5site_1.xvg | awk ′{print $2}’) | \
  column -t > 2dproj_pYsite_+5site_1.xvg
~~~

In principle, projections on these two subvectors can characterize four conformational states given by the two possible states of the pY loop times two possible states of the +5 site (four shaded areas in each panel of Fig. 3B). The two additional states, relative to the α- and β-states characterized above, would be given by *(i)* a closed pY loop with an open +5 site (γ-state) and *(ii)* an open pY loop with a closed +5 site (δ-state) (see Note 5).

As reported in Fig. 3B, a correlation between the pY site and the +5 site is observed in all simulations, where most peptides strictly select for either the α- or for the β-state (Fig. 2B, blue/green areas). Namely, when the pY site is closed, the +5 site is mostly closed (β-state); when the pY site is open, the +5 site is always open (α-state). Consequently, a conformational change at the pY site is accompanied by a concerted conformational change at the +5 site, spanning ∼ 20 Å across the N-SH2 domain (Note 6) [22].

The conformational equilibrium in the N-SH2 domain is determined by a balance of different energetic contributions: *(i)* the intermolecular interactions between the ligand and the N-SH2 domain, and *(ii)* the intramolecular interactions between the residues of N-SH2. This can be rationalized for three typical binding modes using a diagram as reported in Fig. 4:

**Figure 4.**
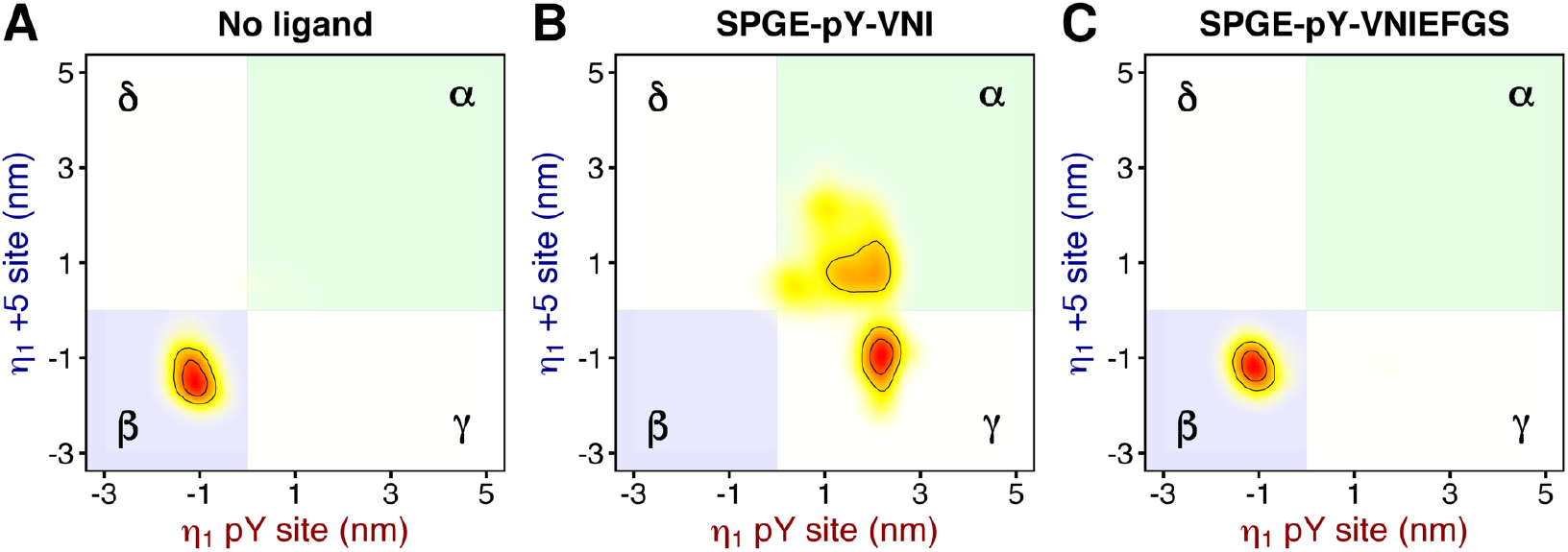
Projection of the N-SH2 trajectories, either (A) unbound N-SH2 or (B–C) bound N-SH2 with peptides of different lengths (subplot titles), projected onto the PCA subvectors of the pY loop (abscissa) and of the +5 site (ordinate). The region corresponding to the α-state (pY loop closed, +5 site closed) is shaded in green, the region of the β-state (pY loop open, +5 site open) is shaded in blue. The regions corresponding to the additional γ-state (pY loop closed, +5 site open) and δ-state (pY loop open, +5 site closed) are reported as white areas.

1. In absence of a ligand (Fig. 4A), the β-state represents the most populated conformational state, presumably favored by the formation of inter-strand H-bonds between the closed central β-sheet.
2. In presence of a truncated ligand, such as SPGEpYVNI (Fig. 4B), which contains a phosphotyrosine but lacks the residues flanking the +5 site, the pY loop–phosphate interactions prevail over inter-strand H-bonds of the central β-sheets, leading to closure of the pY loop and opening of the β-sheets. The α-state is one of the most populated conformations, along with the cognate γ-state. It is worth noting that in absence of residues binding the specificity +5 site, the pY loop is stuck in closed conformation while the conformation of the +5 site is formally decoupled, since it may be either closed (α-state) or open (γ-state). Therefore, the residues binding the specificity site play an essential role in determining the conformation of the N-SH2 domain. This is achieved by their electrostatic attraction or steric repulsion with the flexible loops lining the +5 site.
3. With a specific binding mode of the ligand SPGEpYVNIEFGS, the β-state results the most populated conformation (Fig. 4C). In this specific binding mode, a bulky phenylalanine at position +5 occupies the cleft of the +5 site, thereby keeping the +5 site open. This conformation stabilizes the central β-sheet, aiding the inter-strand H-bonds of the β-sheet to prevail over the pY loop–phosphate H-bonds. Consequently, β-sheets close and pY-loop opens.

### 3.5 Similarities and differences in dynamics between C-SH2 and N-SH2

Can we use the eigenvectors obtained from the cumulative trajectory of the N-SH2 domain to analyze and to compare the conformations adopted by another SH2 domain such as C-SH2? This mostly depends on the structural homology and on the sequence conservation between the two SH2 domains. If possible, it represents a unique chance to disclose similarities and differences between the conformations adopted by the two SH2 domains of SHP2.

Figure 5 presents the sequence alignment after structural superposition of the C-SH2 domain over the N-SH2 domain. The two sequences are characterized by a high sequence conservation. The major difference is found in the length of the loop connecting the βC and βD strands, which is irrelevant for the PCA since it is part neither of the pY loop nor of the +5 site. The pY loop (Ser^34^– Phe^41^ for N-SH2, Ser^140^–Phe^147^ for C-SH2) is well conserved and of the same length in the two SH2 domains (8 a.a.), therefore the corresponding eigenvector may be used without modifications. On the other hand, the +5 site (Gln^57^–Glu^97^ for N-SH2, Arg^173^–Gln^211^ for C-SH2) is less conserved, as expected for a specificity site. The +5 site sequence in C-SH2 is shorter (39 a.a. vs. 41 a.a.) and misses two residues, corresponding to His^85^ and Gly^86^ in N-SH2. Therefore, it is not possible to bijectively map the residues of the +5 site in N-SH2 over the residues of the +5 site in C-SH2. However, all the residues of the +5 site in C-SH2 can be mapped over the corresponding residues in N-SH2, and therefore a comparison is possible only if the residues His^85^ and Gly^86^ are neglected in +5 site definition.

**Figure 5.**
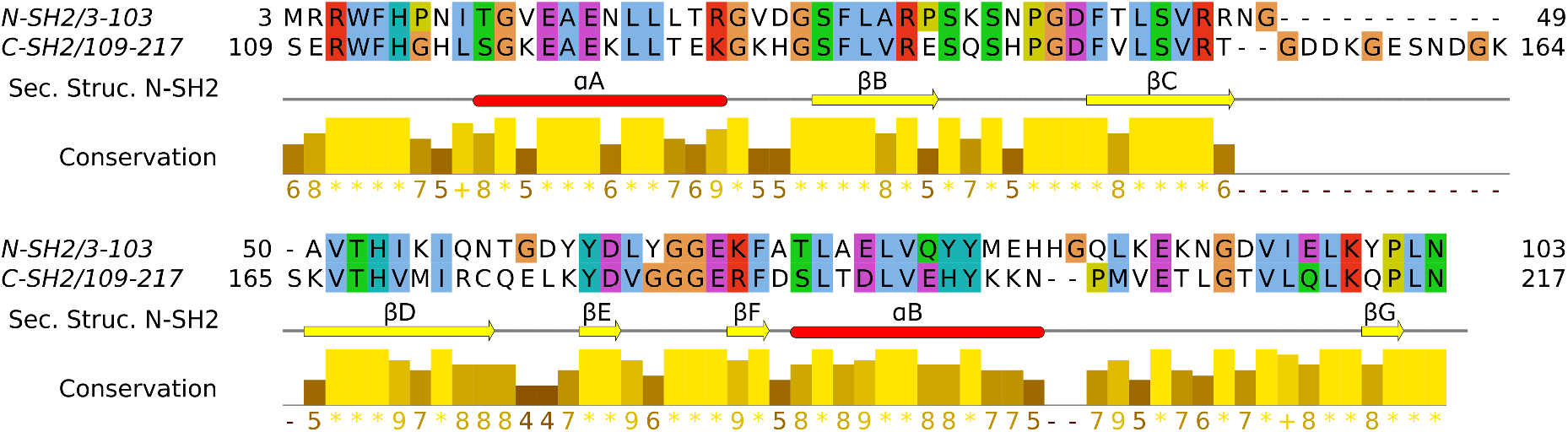
Sequence alignment after structure superposition of the C-SH2 domain structure over the N-SH2 domain structure. The conserved amino acids are highlighted according their properties and the following color scheme: hydrophobic, aliphatic (blue), basic, positively charged (red) acidic, negatively charged (magenta), polar (green), aromatic (cyan), glycines (orange), prolines (yellow). Conservation is visualized on the alignment as a histogram giving the score for each column. The score measures the number of conserved physico-chemical properties conserved for each column of the alignment. Score calculation is derived from the AMAS method of multiple sequence alignment analysis [23]. Fully conserved columns are indicated by “*” (score of 11, grouping all default amino acid properties). Columns with mutations, where all properties are conserved, are marked with a “+” (score of 10, indicating all properties are conserved).

First to proceed with the PCA of C-SH2, create a reference structure of C-SH2, which overlaps to the reference structure of N-SH2 used previously. This task is performed by the *confrms* module that carries out the least-squared fit using two corresponding groups of atoms and does not require that the two structures have the same number of atoms. Therefore, mapping the N-SH2 core, previously used, over C-SH2, obtain the following sequence ranges: Phe^113^–Glu^139^, Asp^146^–Thr^153^, Lys^166^–Cys^174^, Leu^177^–Val^181^, Phe^187^–Tyr^197^, Met^202^–Glu^204^, Val^209^–Pro^215^.

~~~
$ gmx confrms -f1 ref_structure_N-SH2.gro -n1 index_N-SH2   \
  -f2 any_structure_C-SH2.gro -n2 index_C-SH2 -one -o ref_structure_C-SH2.gro
~~~

Then, extract from the filtered cumulative trajectory of N-SH2 the sub-eigenvector for the +5 site according the new definition, which excludes only the residues His^85^ and Gly^86^.

~~~
$ gmx covar -f cumulative_traj_filtered.xtc -s ref_structure_N-SH2.gro   \
  -n index.ndx -o new_eigenvec_5site.trr
~~~

Finally, calculate the projections onto the subvectors for any trajectory of C-SH2 (Note 7):

~~~
$ gmx anaeig -v eigenvec_pYsite.trr -s ref_structure_C-SH2.gro -n index.ndx   \
  -f any_traj_C-SH2.xtc -first 1 -last 1 -proj proj_pYsite.xvg
$ gmx anaeig -v new_eigenvec_+5site.trr -s ref_structure_C-SH2.gro   \
  -n index.ndx -f any_traj_C-SH2.xtc -first 1 -last 1 -proj proj_+5site.xvg
~~~

Again, plot the projection onto the pY site subvector versus the projection onto the +5 site subvector:

~~~
$ paste <(sed ′/^[#@&]/d’ proj_pYsite.xvg | awk ′{print $2}’)   \
        <(sed ′/^[#@&]/d’ proj_+5site.xvg | awk ′{print $2}’) |   \
  column -t > 2dproj_pYsite_+5site.xvg
~~~

In absence of a ligand, the C-SH2 domain adopts a β-like conformation, with both the pY loop and the +5 site in open conformation (Fig. 6A). This is in line with that observed in the N-SH2 domain (Fig. 4A). However, in presence of a full-length phosphopeptide (Fig 6B-F), the C-SH2 domain adopts a γ-like conformation, with the pY loop closed and the +5 site still open. In case of the C- SH2 domain, a coupling between the affinity and the specificity site is not observed, and the region of the domain lining the +5 site does not undergo any significant conformational change to accommodate the phosphopeptide. This behavior is in contrast to our findings for the N-SH2 domain, as discussed above (see Note 8).

**Figure 6.**
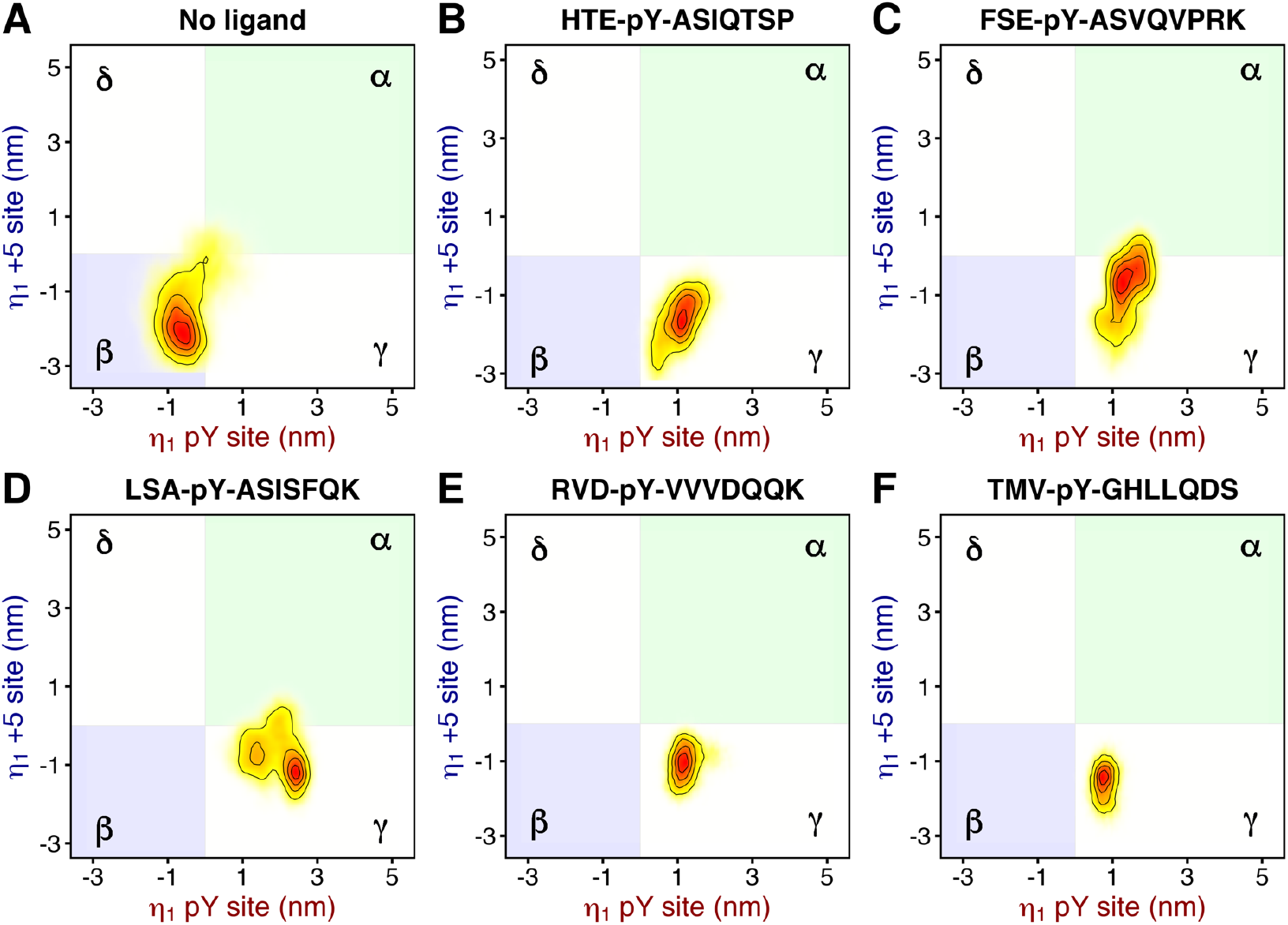
Projection of C-SH2 trajectories, either (A) unbound C-SH2 or (B–F) bound C-SH2 with different peptides (subplot titles) projected onto the PCA subvectors of the pY loop (abscissa) and of the +5 site (ordinate). The region corresponding to the α-state (pY loop closed, +5 site closed) is shaded in green, the region of the β-state (pY loop open, +5 site open) is shaded in blue. The regions corresponding to the additional γ-state (pY loop closed, +5 site open) and δ-state (pY loop open, +5 site closed) are reported as white areas.

## 4. Notes

1. More commonly, principal component analysis is performed on equilibrated trajectories of a biomolecule in a given state (i.e. binding, redox, folding, etc). In that case, the principal modes of motion are representative of the largest structural rearrangements that the biomolecule undergoes during the simulations, and they provide information on the dynamics and the structural plasticity of the biomolecule in a particular condition. Alternatively, the analysis can be performed on a cumulative trajectory encompassing several distinct states of the same biomolecule. Critically, the cumulative trajectory would contain only the part of the system shared among all original trajectories. This time, the principal modes of motion do not describe only the conformational transitions undergone by the biomolecule in a particular state or simulation; instead, they describe the overall conformational space accessible to the biomolecule, as collected from all simulations. Namely, the principal modes of motion consider, besides the dynamics of the system in every state, even the conformational transitions that the biomolecule may undergo for switching from a state to another, even if these transitions were not observed in any individual trajectory. Hence, such analysis reveals conformational transitions between distinct states of a biomolecules, and it indicates how to describe such transitions by molecular structural features, such as interatomic distances or angles, which can be utilized in following analyses or simulations.
2. The trajectory joining the extreme configurations can be used to systematically search the interatomic distances that best linearly correlate with the projection onto the eigenvector.
3. This approach permitts to overcome the problem of poor sampling that affect the individual trajectories, and to detect an allosteric interaction between the affinity site and the specificity site of N-SH2.
4. It is important to perform the superposition using the same reference structure and the same reference group used in the previous covariance matrix calculation on the non-filtered cumulative trajectory. The covariance analyses performed on the filtered trajectory (i.e. projected onto the original full eigenvector) will generate every time only one eigenvector with a non-zero eigenvalue. Any first eigenvector will retain the normalized components of the original full eigenvector, corresponding to the group of atom coordinates used for the analysis.
5. The eigenvector describing a coupling between two sites can be always split into the subvectors representing each site. We can combine these subvectors to generate a symmetric configuration (where the two sites are correlated like in the original eigenvector) and an antisymmetric configuration (where the two sites are anticorrelated) and use the plane of the two subvectors to represent the correlated and anticorrelated motions. Such analysis is, to our knowledge, still rare in the literature. The advantage of this procedure, compared the use of a projection onto a single eigenvector is that we can quickly visualize the component representing the deviation from a perfect correlation between the two sites, which would be otherwise orthogonal to the original eigenvector.
6. Critically, it is unlikely that individual simulations explore all possible binding modes of a phosphopeptide within 1 µs of simulation time because the interconversion between binding modes would first involve a weakening of peptide–N-SH2 interactions, before the required side chain and backbone reorientations could occur. Therefore, each individual simulation likely showed only a partial view on the accessible binding modes of each peptide, and probably in every simulation the N-SH2 domain is mostly stuck in one of the possible conformations. Consequently, such equilibrium MD simulations, taken individually, are insufficient for deciding whether a specific peptide sequence selects for the α- or the β-state and for estimating an equilibrium constant between the two conformational states. However, the individual simulations taken together are representative of the N-SH2 domain in bound form to reveal a correlation between the affinity and the specificity site.
7. Since the subvectors were obtained from the eigenvector of N-SH2, they are not strictly representative of the collective modes of motion in C-SH2. However, this procedure permits a direct comparison in order to test if the collective motions in N-SH2 are conserved in C- SH2, thereby revealing similarities and differences between the N-SH2 and C-SH2 dynamics.
8. This application confirms that PCA can be used to compare dynamics between homologous domains, not just SH2, evidencing similarities and differences.

